# Anti-tubercular potential and pH-driven mode of action of salicylic acid derivatives

**DOI:** 10.1101/2024.09.22.614344

**Authors:** Janïs Laudouze, Thomas Francis, Emma Forest, Frédérique Mies, Jean-Michel Bolla, Céline Crauste, Stéphane Canaan, Vadim Shlyonsky, Pierre Santucci, Jean-François Cavalier

## Abstract

In the search for new anti-tuberculosis drugs with novel mechanisms of action, we evaluated the antimycobacterial activity of a panel of eight phenolic acids against four pathogenic mycobacterial model species, including *M. tuberculosis*. We demonstrated that salicylic acid (**SA**), as well as the iodinated derivatives 5-iodo-salicylic acid (**5ISA**) and 3,5-diiodo-salicylic acid (**3,5diISA**), displayed promising antitubercular activities. Remarkably, using a genetically encoded mycobacterial intrabacterial pH reporter, we describe for the first time that **SA, 5ISA, 3,5diISA** and the anti-inflammatory drug aspirin (**ASP**) act by disrupting the intrabacterial pH homeostasis of *M. tuberculosis* in a dose-dependent manner under *in vitro* conditions mimicking the endolysosomal pH of macrophages. In contrast, the structurally related second-line anti-TB drug 4-aminosalicylic acid (**PAS**) had no pH-dependent activity and was strongly antagonized by L-methionine supplementation, thereby suggesting distinct modes of action. Finally, we propose that **SA, ASP** and its two iodinated derivatives could restrict *M. tuberculosis* growth in a pH-dependent manner by acidifying the cytosol of the bacilli; therefore, making such compounds very attractive for further development.

## Introduction

Tuberculosis (TB), caused by its aetiologic agent *Mycobacterium tuberculosis* (*M. tuberculosis*), remains a global public health issue worldwide. The latest World Health Organization report estimates that in 2022, 1.3 million people died from TB and approximately 10.6 million people contracted the disease.^[1]^

The current standard drug regimen, composed of isoniazid (INH), rifampin (RIF), ethambutol (EMB) and pyrazinamide (PZA), is generally effective with an estimated average success rate of 88% on TB-susceptible cases.^[1]^ However, with four distinct drugs to be taken for at least 6 months, this lengthy and toxic regimen often affects compliance, leading to treatment failures and the emergence of drug-resistant strains.^[1]^ Indeed, recent observations by the World Health Organization suggest that the number of drug-resistant cases (e.g. RR-TB; MDR-TB; XDR-TB) is increasing in many areas of the world, leaving clinicians with very few successful therapeutic interventions.^[1]^ In that context, the discovery and/or the development of chemical entities that can kill *M. tuberculosis* more efficiently is urgently needed to better control the global TB pandemic.^[2]^

Phenolic acids (PA), also known as phenol carboxylic acids, are a specific class of aromatic compounds that contain both a phenolic ring and a carboxyl functional group. This class of compounds represents the simplest polyphenols in terms of chemical structure, and some PA have been described as very promising antimicrobial agents.^[3-5]^ Because of their antibacterial, antiviral and antifungal properties as well as low background toxicity towards host cells, PA and their derivatives have found widespread applications as preservatives in food, pharmaceutical, and cosmetic products.^[6]^ In addition to these properties, PA have been also described for other health protective effects such as anti-oxidants and anti-inflammatory, therefore constituting a promising class of compounds for therapeutic applications.^[7]^

In the context of TB, several PA have been identified with good anti-TB properties^[5, 8-11]^ and thus proposed as potential candidates in TB therapy.^[5]^ Historically, salicylic acid (**SA**) and its analogue, the 4-aminosalicylic acid also known as *para*-aminosalicylic acid (**PAS**) have been identified as potential antitubercular in the 1940’s.^[12-15]^ As a direct consequence, **PAS** was then introduced in anti-TB therapies and has been proven to be effective to cure TB when administered alone or co-administered with streptomycin to prevent resistance emergence.^[9, 13]^ This was before the development of the current standard anti-TB regimen; nevertheless, **PAS** remains in the anti-TB arsenal and continues to be used nowadays as second-/third-line drug in some cases.^[16]^

Recently, Zhang and colleagues reported that *M. tuberculosis* was highly susceptible to a wide range of weak acids, including several PA compounds such as benzoic acid (**BA**), **SA** or the anti-inflammatory drug acetyl-salicylic acid known as aspirin (**ASP**).^[9]^ Complementary studies focusing on other pathogenic strains have also reported that **SA** and **ASP** could be used alone or in combination with known antibiotics to enhance antimicrobial susceptibility using *in vitro* and *in vivo* biological systems.^[17-19]^ However, whether such strategy could be applied to *M. tuberculosis* with established molecules and/or new analogs has not been thoroughly investigated.

In that context, we sought to test the antibacterial activity of a panel of PA, including **SA** derivatives, against four pathogenic mycobacterial model species, including *M. tuberculosis*. Using this approach, we demonstrated that parental **SA**, as well as some iodinated derivatives, display promising antitubercular activities. Based on their chemical structures and their well-established weak acid properties, we have tested the pH-dependency of **SA** and its derivatives and further established that some of them exhibit significant pH-driven activities. By using a genetically encoded mycobacterial intrabacterial pH reporter, we describe for the first time that **SA**, iodinated **SA**-derivatives and the anti-inflammatory drug **ASP** are able to disrupt the intrabacterial pH of *M. tuberculosis* in a dose-dependent manner under *in vitro* conditions mimicking the endolysosomal pH of macrophages. Conversely, the structurally related second-line anti-TB drug **PAS** revealed different inhibitory profiles, a strong antagonism with L-methionine supplementation and no pH-dependency suggesting distinct modes of action. Finally, we propose a model that could explain how **SA** and some non-cytotoxic **SA** derivatives restrict *M. tuberculosis* growth in a pH-dependent manner by acidifying its cytosol; therefore, making such compounds very attractive for further development.

## Results and Discussion

### Screening and identification of inhibitory properties of a small subset of PA on mycobacterial model species

We recently described the antibacterial activity of a subset of PA and iodinated-PA on the Gram^(+)^ opportunistic pathogen *Staphylococcus aureus*.^[20]^ Results showed that although native PA did not impact growth at concentration up to 1 mM (*i.e*., 140-200μg/mL), some of their iodinated derivatives displayed better inhibitory activities with minimal inhibitory concentration (MIC) values comprised around 400-700μM (*i.e*., 120-180μg/mL).^[20]^

Based on these observations, we have conducted a more comprehensive susceptibility testing investigation of both parental and iodinated forms of PA against the Gram^(-)^ bacteria *E. coli* and *P. aeruginosa*; and four mycobacterial species, which include the saprophytic species *M. smegmatis*, the opportunistic pathogens *M. marinum* and *M. abscessus* and finally the tubercle bacilli, *M. tuberculosis*.

The chemical structure of the eight compounds tested, gallic acid (**GA**), 2-iodo-gallic acid (**IGA**), *trans*-ferulic acid (***t*FA**), 2-iodo-ferulic acid (**2IFA**), salicylic acid (**SA**), 5-iodo-salicylic acid (**5ISA**), 3,5-diiodo-salicylic acid (**3,5diISA**) and 5-iodo-2-methoxy benzoic acid (**2M,5IBA**) are displayed in **Scheme 1**, and their inhibitory parameters obtained by performing susceptibility testing are reported in **Table 1**.

**Table 1.**
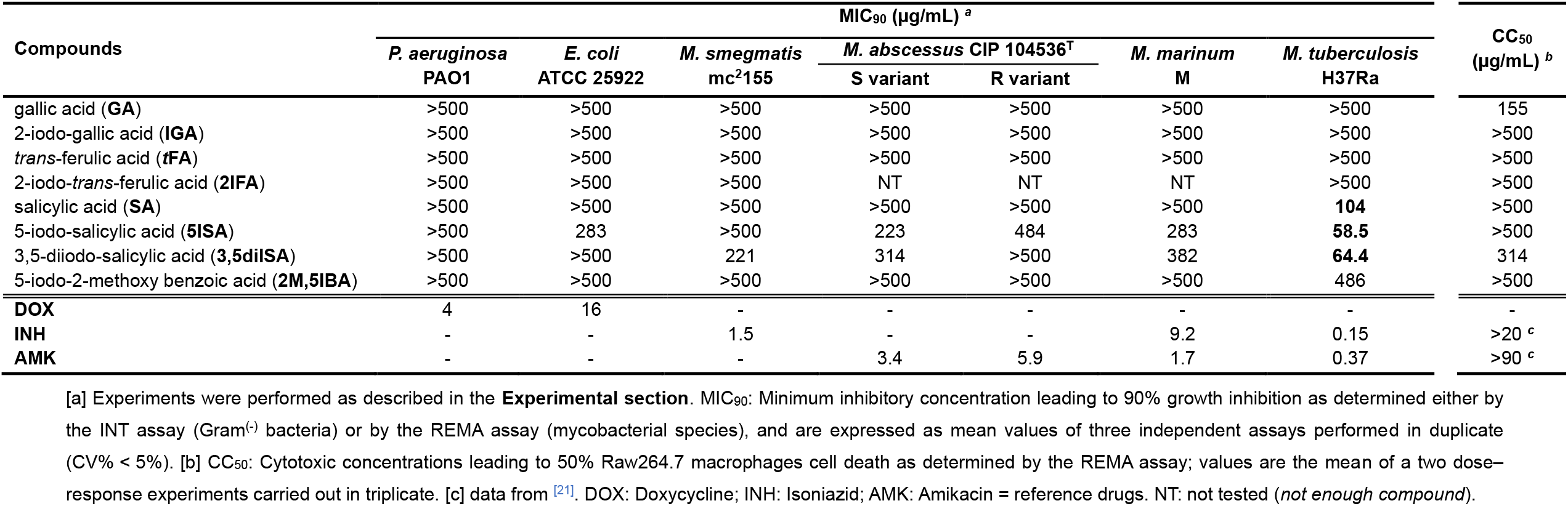
Antibacterial activities of the eight phenolic compounds compared to standard antimicrobial agents against two Gram^(-)^ bacteria and four mycobacterial strains.

**Scheme 1.**
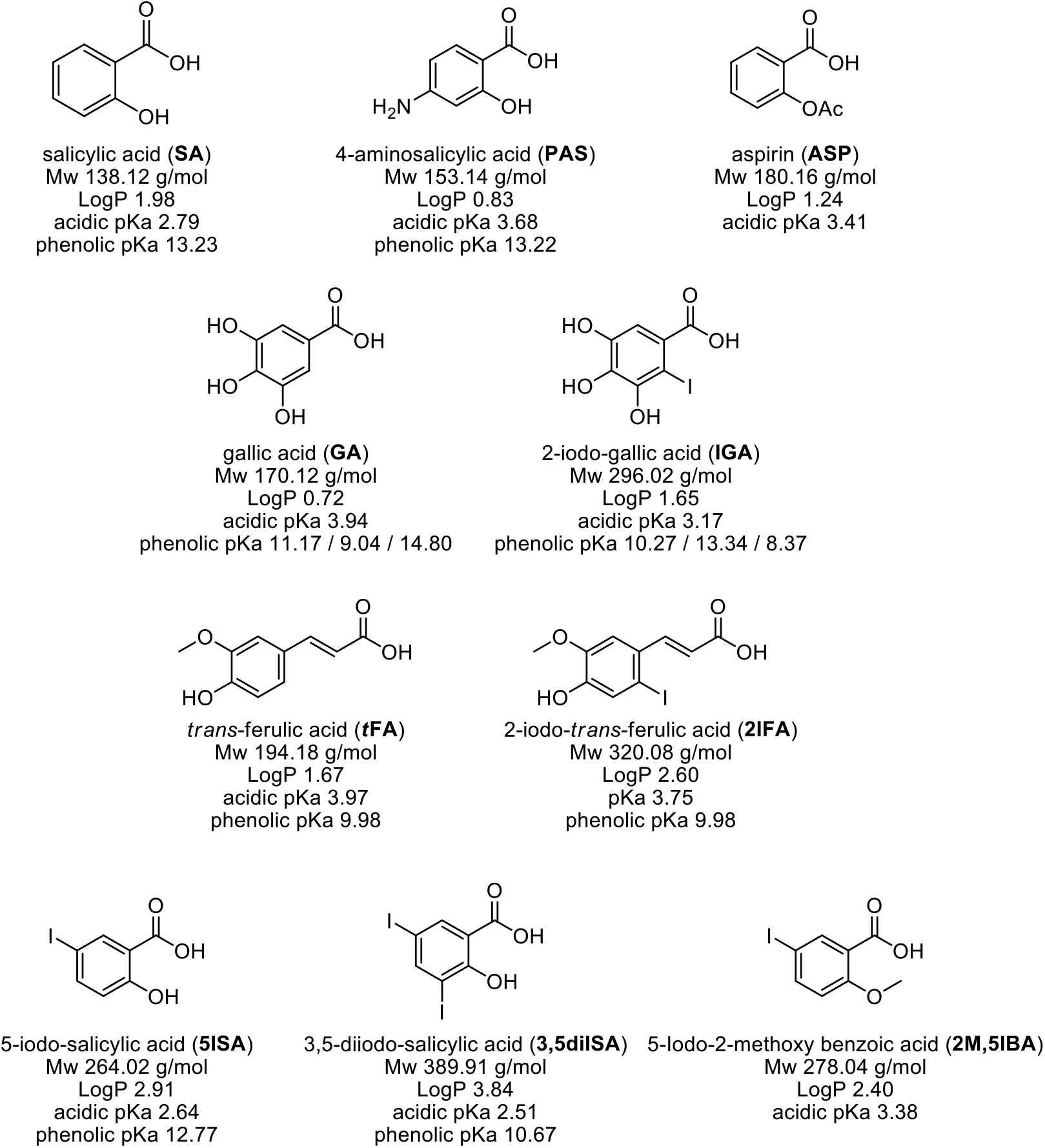
Structure and physico-chemical properties of the PA compounds investigated in this study.

First, susceptibility testing against the two Gram^(-)^ bacterial strains, *E. coli* and

*P. aeruginosa*, showed that the eight molecules were poorly active against the two strains with MIC_90_ values ranging from 283μg/mL to higher than 500μg/mL (**Table 1**).

Further investigation and analysis of fitted growth inhibition profiles and MIC_90_ values against mycobacteria demonstrated that **GA, IGA, *t*FA** and **2IFA** compounds were almost completely inactive against the species tested with MIC_90_ values superior to 500μg/mL (**Table 1**). On the other hand, SA and its derivatives displayed heterogeneous growth inhibitory activities across the various species with MIC_90_ ranging from 58.5 and up to 486μg/mL (**Table 1**). Of importance, *M. tuberculosis* was found to be more susceptible to iodinated SA-derivatives than *M. smegmatis, M. abscessus* S and R morphotypes and *M. marinum*. In fact, **5ISA** and **3,5diISA** strongly inhibited the growth of *M. tuberculosis* with relatively good MIC_90_ values of 58.5 and 64.4μg/mL, respectively, which are 3.4-8.3 times lower than those obtained for the other mycobacterial strains. As previously observed with *S. aureus*,^[20]^ addition of iodine atoms slightly increased the antibacterial activity of the resulting iodinated derivatives *versus* the parental **SA** molecule (MIC_90_ = 104μg/mL) against *M. tuberculosis*.

Finally, the toxicity of the eight PA against murine Raw264.7 macrophages was investigated using a classical dose–response assay.^[22]^ The calculated response parameter was the CC_50_, which corresponds to the concentration required to elicit a 50% decrease in cell viability compared to the control (**Table 1**). Except for **GA** (CC_50_ = 155±16μg/mL) and **3,5diISA** (CC_50_ = 314±18μg/mL) for which very moderate levels of toxicity were found, the other compounds were not toxic to Raw264.7 cells, as demonstrated by CC_50_ values > 500 μg/mL. This finding is in line with previous results showing that these compounds do not affect the viability of L929, HeLa, T84 and Caco-2 cells.^[20]^

Given all these findings, amongst the 8 tested phenolic compounds, **SA** and its two iodinated derivatives, **5ISA** and **3,5diISA**, were displaying the best combination of antibacterial activities against *M. tuberculosis* and cytotoxic properties, and were then selected for further mechanistic investigations.

### Analysis of pH-driven activities of SA and SA derivatives

Previous report demonstrated that **SA** and **ASP** antibacterial activities against *M. tuberculosis* were moderate. However, similarly to many weak acids, both compounds displayed interesting pH-driven potency against *M. tuberculosis*.^[9]^ Indeed, according to their chemical properties and the Henderson-Hasselbalch equation, a decrease in extra-bacterial pH should increase the proportion of neutral protonated weak acid [AH], whereas an increase in pH would favor the formation of its negatively charged anion [A^-^], as described for the anti-TB drug pyrazinoic acid.^[9, 23]^ Thus, at acidic pH, the increase in the protonated neutral form might have two major effects that could explain a more potent activity. Firstly, protonation of weak acids to their neutral form has been shown to facilitate their ability to cross biological membranes,^[24-26]^ thereby increasing their intrabacterial concentrations. Secondly, according to this weak acid model, protonated forms may have the ability to unilaterally translocate protons through the mycobacterial envelope by passive diffusion in a pH-dependent manner and, based on the pKa of the carboxylic acid function, subsequently release protons in the pH-neutral bacterial cytosol.^[9, 23-24]^

In that context, we explored whether **SA**, its two iodinated derivatives, **5ISA** and **3,5diISA** as well as the FDA-approved references drugs **PAS** and **ASP** might share a similar primary mode of action and commonly display pH-driven activities. To achieve this goal, we first assessed antibacterial activity in standard 7H9 medium at pH 6.8 and in MES-adjusted pH 5.5 medium.^[27]^ Since Tween-80 has been shown to impact viability at low pH, this detergent was replaced in both media by tyloxapol, a non-hydrolysable non-toxic dispersible agent.^[28]^ The results obtained in 7H9 medium at nearly neutral pH 6.8 in this slightly different methodological system confirmed those obtained above during the multi-species screening, with the **SA** derivatives showing MIC_90_ values in the range of 100-200μg/mL (**Table 2**).

**Table 2.**
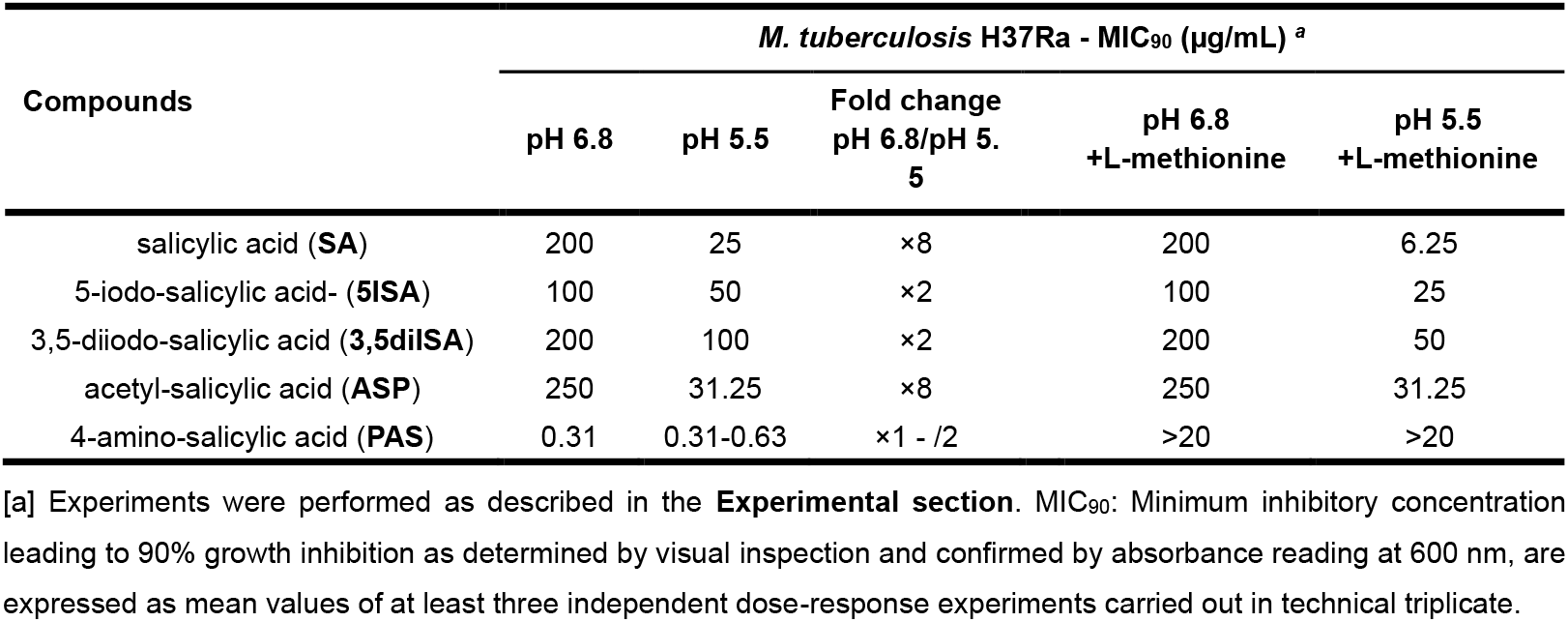
Antibacterial activities of **SA** derivatives on *M. tuberculosis* at near-neutral (pH 6.8) and acidic (pH 5.5) pH in the presence or absence of the **PAS** antagonist methionine.

| Compounds | *M. tuberculosis* H37Ra - MIC<sub>90</sub> (µg/mL) <sup>a</sup> |  |  |  |  |
| --- | --- | --- | --- | --- | --- |
|  | pH 6.8 | pH 5.5 | Fold change<br>pH 6.8/pH 5.5 | pH 6.8<br>+L-methionine | pH 5.5<br>+L-methionine |
| salicylic acid (**SA**) | 200 | 25 | ×8 | 200 | 6.25 |
| 5-iodo-salicylic acid- (**5ISA**) | 100 | 50 | ×2 | 100 | 25 |
| 3,5-diiodo-salicylic acid (**3,5diISA**) | 200 | 100 | ×2 | 200 | 50 |
| acetyl-salicylic acid (**ASP**) | 250 | 31.25 | ×8 | 250 | 31.25 |
| 4-amino-salicylic acid (**PAS**) | 0.31 | 0.31-0.63 | ×1 - /2 | >20 | >20 |

[a] Experiments were performed as described in the **Experimental section**. MIC<sub>90</sub>: Minimum inhibitory concentration leading to 90% growth inhibition as determined by visual inspection and confirmed by absorbance reading at 600 nm, are expressed as mean values of at least three independent dose-response experiments carried out in technical triplicate.

In contrast, performing the experiments at acidic pH 5.5 revealed a strong compound-dependent acidic pH-mediated potentiation, with changes in MIC_90_ from 2-to 8-fold (**Table 2**). The best effect was observed with the reference compound **SA** with an 8-fold decrease in MIC_90_, shifting from 200 to 25 μg/mL. This latter result is in perfect agreement with previously published observations from Zhang *et al*. who reported a better anti-TB activity of **SA** at acidic pH 5.5 (MIC_90_ = 10-20 μg/mL) than at neutral pH 6.8 (MIC_90_ of 50-100 μg/mL).^[9]^ Determination of the MIC_90_ of **ASP** showed a very similar pattern, with MIC_90_ values at pH 6.8 at 250 μg/mL and 31.25 μg/mL when assessed at pH 5.5 resulting in a an 8-fold decrease.

Regarding the two iodinated derivatives, **5ISA** and **3,5diISA**, a twofold decrease in MIC_90_ values was observed at pH 5.5 *vs*. pH 6.8 (**Table 2**). Analysis of the calculated pKa of each compound (**Scheme 1**) failed to establish a clear quantitative link between the observed pH dependence and the ability of the COOH function to gain protons,^[9]^ suggesting that more complex or multiple parameters may influence the activities of these compounds.

As expected, the FDA-approved anti-TB drug **PAS** displayed the best activity with MIC_90_ value of 0.31μg/mL when assessed at pH 6.8 and was not significantly impacted by acidic pH, suggesting that **PAS** does not act as a pH-dependent compound (**Table 2**).

Altogether, our results confirm that **SA** and **ASP** exhibit strong pH-dependent potentiation,^[9]^ whereas iodinated analogues are only slightly affected by this parameter. This was also observed for **PAS** which consistently inhibited bacterial growth regardless of environmental pH. Such findings confirm that structural analogues, with very minor differences can still exhibit different pH-dependent inhibitory activity.

### Methionine-mediated antagonism occurs for the folate biosynthesis targeting compound PAS but not any other SA derivatives

Recent studies have demonstrated that L-methionine supplementation can drastically affect the antibacterial activity of **PAS** and other anti-folate drugs.^[29-30]^ To gain further insight onto the mode of action of **SA** and its derivatives, we tested whether L-methionine could also have a negative impact on their inhibitory features at both near-neutral and acidic pH. First, and as expected, supplementation with 10 μg/mL of L-methionine triggered an important change in **PAS** inhibitory activity with MIC_90_ shifting from 0.3125 μg/mL to > 20 μg/mL at both pH 6.8 or pH 5.5 (**Table 2**). This 64-fold increase in MIC_90_ is in perfect agreement with previously published observations from Howe *et al*.^[30]^ In contrast, in the presence of L-methionine at both pH, identical or a twofold decrease in MIC_90_ values were observed for **ASP, 5ISA** and **3,5diISA** (**Table 2**). Surprisingly, L-methionine supplementation potentiated the activity of **SA** by 4-fold at acidic pH but not at near-neutral pH (**Table 2**). This result was not really expected, but the beneficial effect of L-methionine has been already reported in the literature.^[31]^ Indeed, L-methionine has been shown to potentiate several classes of antibiotics including macrolides, cyclines or fluoroquinolones, and such process has been proposed to be associated with a decrease in efflux and/or alteration of the oxidative stress.^[31]^

Taken together, our results suggest that the mode of action of **SA, 5ISA, 3,5diISA** and **ASP** may be clearly distinct from that of **PAS** which is strongly antagonized by L-methionine. Accordingly, we propose that the *M. tuberculosis* folate pathway is unlikely to be targeted by **SA, 5ISA, 3,5diISA** and **ASP**.

### SA and SA derivatives acidify *M. tuberculosis* cytosol in a dose-dependent and pH-dependent manner

We capitalized from a well-established genetically encoded intrabacterial pH (IBpH) reporter (**Figure 1**)^[28, 32-33]^ to assess if **SA** and **SA**-iodinated derivatives would be able to acidify IBpH as previously hypothesized in the case of weak acids.^[9]^

**Figure 1.**
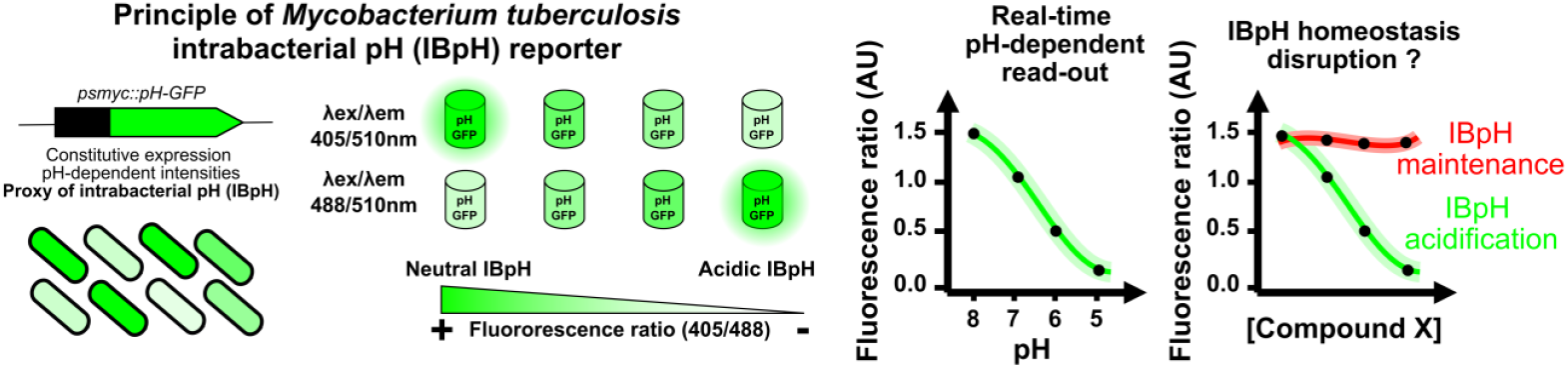
Schematic representation of *M. tuberculosis* intrabacterial pH (IBpH) homeostasis monitoring in real time using the genetically-encoded pH-GFP sensitive ratiometric reporter. The *pH-GFP* gene is encoded by the pUV15-pHGFP episomal vector and its expression is driven under the strong constitutive promoter *psmyc*. The dual pH-GFP excitation/emission wavelengths are inversely responding to pH exposure, thereby displaying distinct fluorescence intensity profiles which can be monitored noninvasively overtime. Analysis can be expressed as fluorescence intensity ratio that are obtained by dividing the fluorescence intensity acquired with excitation/emission channels of 405/510 nm by the one at 488/510 nm. A decrease in the 405/488nm ratio highlights the acidification of the bacterial cytosol. A representation of such decrease in ratiometric signal quantification as function of pH is displayed. Finally, this tool can be used to monitor the effect of drugs onto *M. tuberculosis* IBpH homeostasis as highlighted on the far-right panel.

Dose-response analysis performed at pH 6.8 showed that only very high-concentration (50-200 μg/mL) of **SA** triggers IBpH homeostasis perturbation (**Figure 2A – *left panel***). Indeed, such process was only visible at concentrations close to **SA**’s MIC_90_ value with statistically significant changes noticeable when pulsed with **SA** concentrations comprised between 50 and 200 μg/mL (*p*-value<0.05 and *p*-value<0.001, respectively). Increasing protons availability in the extracellular medium by adjusting the pH at 5.5 significantly affected the ability of **SA** to disrupt IBpH homeostasis, as concentrations from 12.5 μg/mL significantly impacted the fluorescence ratio thus highlighting an acidification of the bacterial cytosol (**Figure 2A – *middle panel***). Dose response curve analysis and 4-parameter regression allow us to estimate **SA**’s EC_50_ for IBpH homeostasis disruption. As shown in **Figure 2A** (***right panel***) acidic pH potentiated the effect of **SA** on IBpH, with a drop in EC_50_ of approximately 2.5-fold (38.92 ± 26.40 μg/mL *vs*. 15.05 ± 1.70 μg/mL). This confirms the seminal observation of Zhang and colleagues who reported that **SA** possesses a pH-driven, pH-disruptive effect against the tubercle bacilli.^[9]^

**Figure 2.**
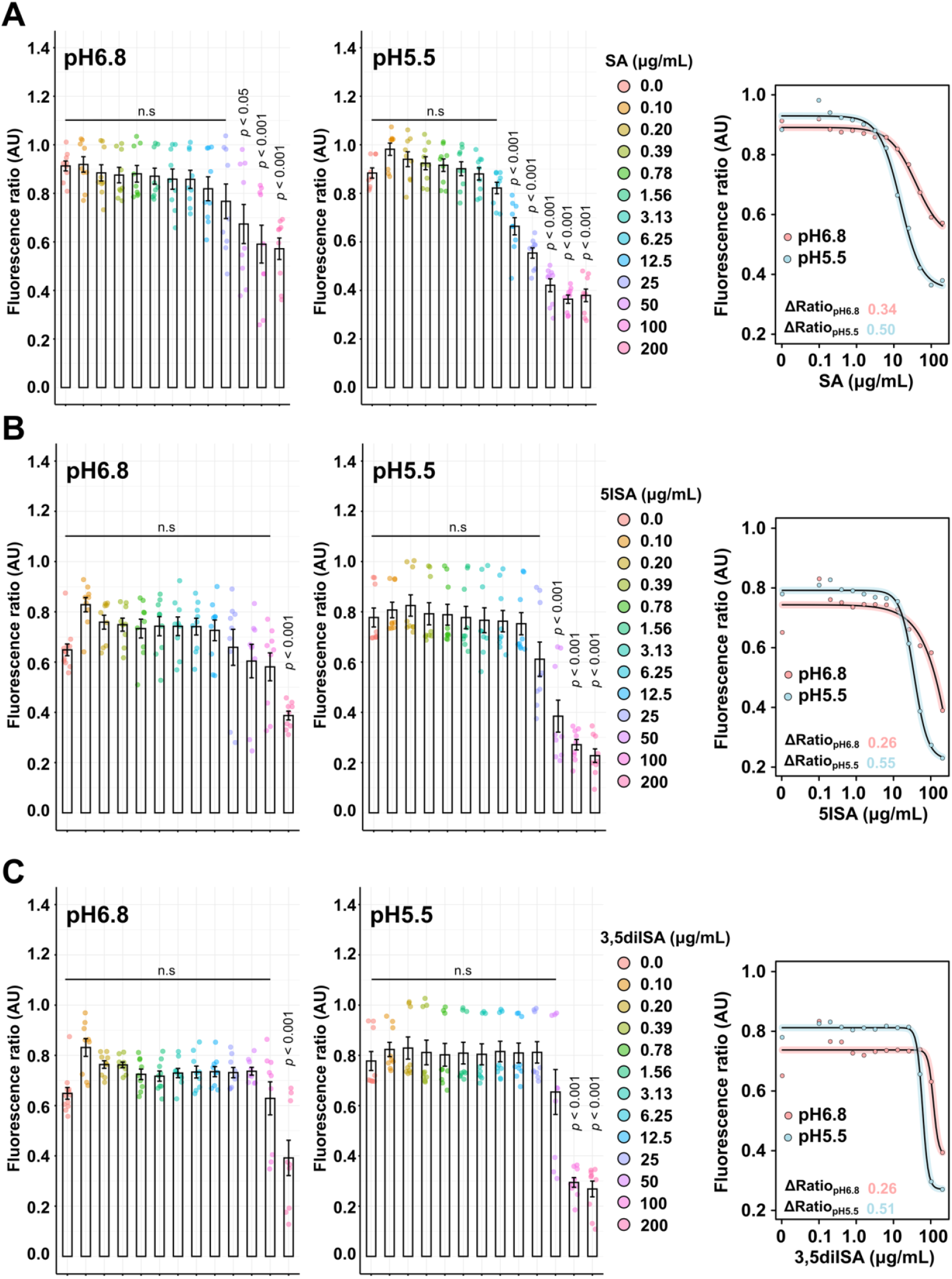
*M. tuberculosis* intrabacterial pH homeostasis disruption by SA and derivatives. **(A-C)** Quantification of *M. tuberculosis* pH-GFP ratio (405/488 nm) in the presence of increasing concentrations of **(A) SA, (B) 5ISA** and **(C) 3,5diISA** for 24 hours. Ratiometric signals were obtained by dividing the fluorescence intensity acquired with excitation/emission channels of 405/510 nm by the one obtained at 488/510 nm. Results from **SA, 5ISA** and **3,5diISA** are displayed from top to bottom respectively. Analysis performed at pH 6.8 are displayed on the left panels whereas analysis performed at pH 5.5 are displayed on the middle panels. The right panels correspond to the 4-parameter nonlinear logistic regression of the data displayed in the left and middle panels respectively. Results were obtained from *n* = 3 biologically independent experiments and are displayed as mean ± SEM. Statistical significance was assessed by comparing the means of each concentration with its respective control condition using one-way ANOVA followed with Tukey’s multiple comparisons test. All *p*-values were considered significant when *p*-value<0.05. EC_50_ determination was not applicable for the compound **5ISA** at pH 6.8 due to a lack of complete sigmoidal response.

The same approach was conducted with the **5ISA** and **3,5diISA** compounds (**Figure 2B-C**). At pH 6.8 a significant perturbation of IBpH homeostasis was only visible at the highest concentration tested (200 μg/mL – *p*-value<0.001) (**Figure 2B-C – *left panels***). At pH 5.5, both derivatives disrupt IBpH homeostasis similarly to **SA**. This shift in *M. tuberculosis* iBpH was observed at lower concentrations of 50 μg/mL and 100 μg/mL for **5ISA** and **3,5diISA**, respectively (*p*-value<0.001) (**Figure 2B-C – *middle panels***). Similar trend was observed on their corresponding fitted graphs, highlighting a pH-disruptive effect (**Figure 2B-C – *right panels***).

The analysis of the difference between the maximum and minimum fluorescence ratio values (expressed as ΔRatio_pH6.8_ or ΔRatio_pH5.5_) showed that pH 5.5 induced a greater amplitude in IBpH difference for the same concentration of compounds. With **SA**, the ΔRatio_pH6.8_ and ΔRatio_pH5.5_ were 0.34 and 0.50, respectively, therefore indicating a greater cytosolic acidification when more extra-bacterial protons are available (**Figure 2A – *right panel***). Almost identical values to **SA** were obtained with **5ISA** and **3,5diISA**, with ΔRatio_pH5.5_ of 0.55 and 0.51, respectively.

Overall, all these results confirm that the introduction of iodinated groups on the parent **SA** molecule had only a minor effect on its antibacterial activity. On the other hand, they retain their ability to be potentiated at acidic pH, and exhibit IBpH disruptive activities closely correlated with their ability to block bacterial growth.

### ASP, but not PAS, displays IBpH disruptive activity reflecting different modes of action

Finally, we tested the ability of the clinically relevant drugs, **PAS** and **ASP**, to disrupt *M. tuberculosis* IBpH (**Figure 3A-B**). Structurally, **ASP** is a very close analogue of **SA** in which the hydrogen that is attached to the phenolic hydroxy group has been replaced by an acetyl group. With a strong pH-dependent potency, we hypothesized that **ASP** may display an important IBpH disruptive activity similar to SA. The dose-response analysis performed showed that ASP had modest but significant effects on IBpH when tested at pH 6.8 and concentrations ranging from 62.5 to 250 μg/mL (*p*-value<0.05 and *p*-value<0.001, respectively) (**Figure 3A**). When the experiment was carried out at acidic pH, **ASP** was able to significantly disrupt IBpH from 31.25 μg/mL, supporting its previously observed pH-dependent activity. Important changes were also observed at lower concentration such as 15.6 μg/mL but failed to reach statistical significance with a *p*-value of 0.06. Determination of ΔRatio_pH6.8_ or ΔRatio_pH5.5_ confirmed this strong pH-dependency as well as a very potent activity on IBpH with values of 0.26 and 0.62, respectively (**Figure 3A**).

**Figure 3.**
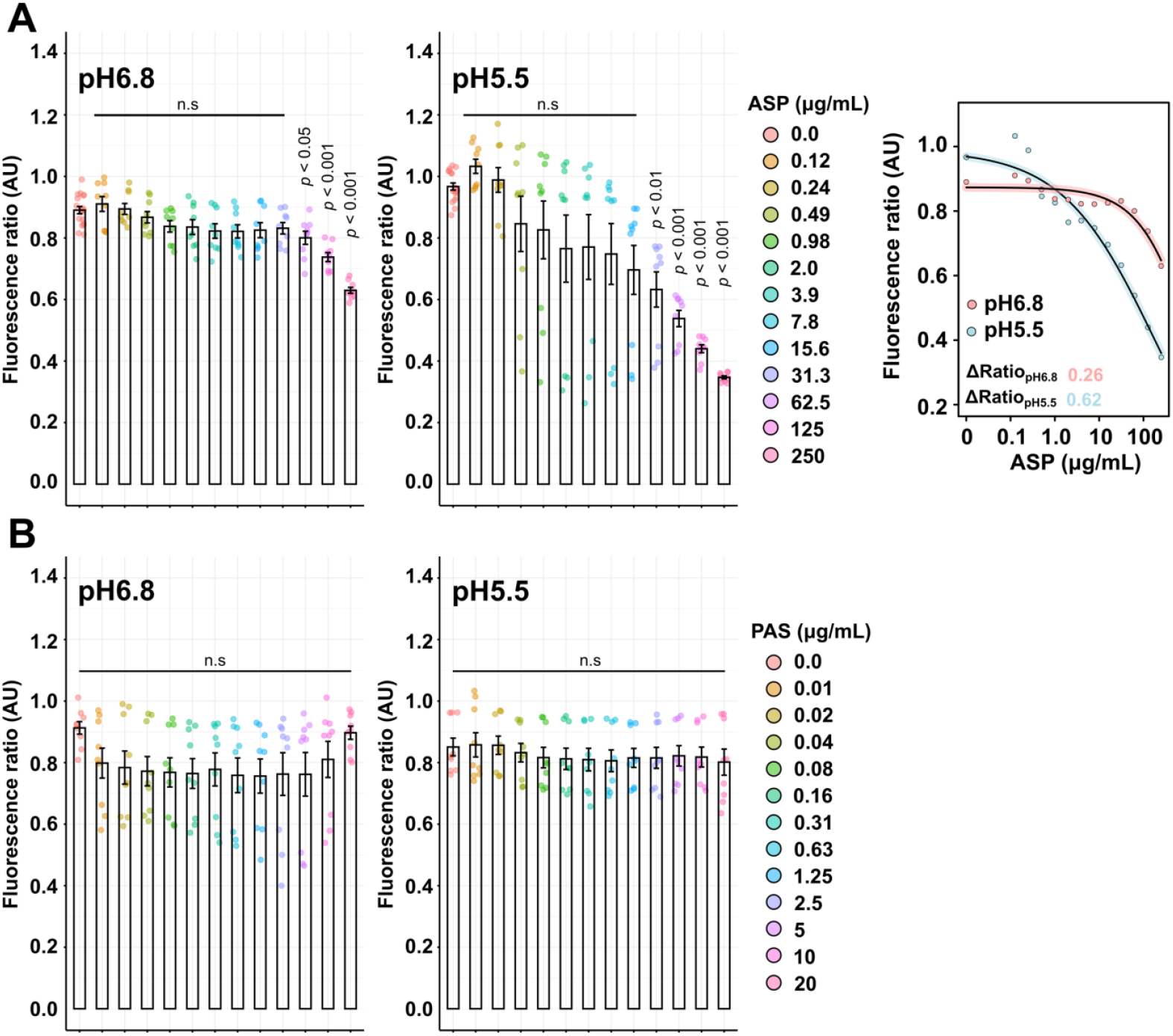
Divergent effects of the clinically available drug ASP and PAS on *M. tuberculosis* intrabacterial pH homeostasis supports different antibacterial mode of action. **(A-B)** Quantification of *M. tuberculosis* pH-GFP ratio (405/488nm) in the presence of increasing concentrations of **ASP (A)** or **PAS *(B)*** for 24 hours. Ratiometric signals were obtained by dividing the fluorescence intensity acquired with excitation/emission channels of 405/510 nm by the one obtained at 488/510 nm. Results from **ASP** and **PAS** are displayed from top to bottom respectively. Analysis performed at pH 6.8 are displayed on the left panels whereas analysis performed at pH 5.5 are displayed on the middle panels. Results were obtained from n = 3 biologically independent experiments and are displayed as mean ± SEM. Statistical significance was assessed by comparing the means of each concentration with its respective control condition using one-way ANOVA followed with Tukey’s multiple comparisons test. All *p*-values were considered significant when *p*-value< 0.05. The right panel of **ASP** corresponds to 4-parameter nonlinear logistic regression of the data displayed in the left and middle panels respectively. No curve-fitting was performed for **PAS** due to a lack of convergence when performing the 4-parameter nonlinear logistic regression.

On the contrary, **PAS** has no effect on IBpH at both pH 6.8 and pH 5.5, even when tested at 20 μg/mL, a concentration 64 times higher than its MIC_90_ (**Figure 3B**). These results further indicate that the anti-tubercular activity of **PAS** does not definitively result from a major modification of the IBpH, but is likely due to the inhibition of folate biosynthesis by targeting the dihydrofolate reductase DfrA,^[34]^ as supported by our own observations of L-methionine-mediated antagonism (**Table 2**).

### Inability to isolate SA and SA-derivatives spontaneous resistant mutants

Previous studies have reported that isolation of spontaneous resistant to weak acids is extremely complicated, likely due to their unique mode of action disrupting intrabacterial pH homeostasis independently of a specific protein target. In that context, Zhang *et al*. suggested a potential highly conserved mechanism by which intracytoplasmic protons alter numerous processes simultaneously and therefore do not allow the selection of spontaneous resistant mutants.^[9]^

To further characterize the mode of action of our compounds, we performed a similar subset of experiments and tried to isolate spontaneous resistant mutants as previously described.^[9, 35]^ For that, 10^6^ CFU were plated on 7H11 agar plates at pH 6.8 containing a concentration corresponding to the MIC, 2×MIC or 5×MIC of compounds. After 4-6 weeks of incubation, no resistant colonies had appeared for the compounds that displayed pH-driven, pH-disruptive activities, suggesting that the isolation of resistant mutants is indeed complex and might require long-term exposure at sub-optimal concentration to select and enrich for mutations that increase fitness in the presence of weak acids as previously demonstrated.^[36]^

Overall, these results do not necessary rule out that these compounds do not have specific target(s) in *M. tuberculosis*, but they are in favor of a potential pH-dependent pH-disruptive activity that precludes the selection of spontaneous resistant as previously suggested by Zhang *et al*. ^[9]^

## Conclusion

In this study, we identify and characterize the pH-dependent and IBpH-disruptive activities of compounds that are derived from **SA**, a very simple PA. We demonstrate that iodination of **SA** does not alter the antibacterial activity of the resulting **5ISA** and **3,5diISA** derivatives, and could constitute an interesting way to synthetize new antibacterial molecules. In addition, our mechanistic studies reveal that the anti-inflammatory drug **ASP** is also a potent anti-TB drug that is potentialized at acidic pH and further acidify *M. tuberculosis* cytosol. This mode of action appears to be different from that of the well-established anti-TB drug **PAS**, which is also an analogue of **SA**. Altogether, these observations suggest that PA and more precisely **SA**-derivatives may represent an interesting subset of molecules that could open new avenues for the development of new antibacterial entities with more potent activity in physiologically relevant microenvironments encountered by *M. tuberculosis*.

## Experimental section

### Chemical compounds and computed properties

Compounds Gallic acid (**GA**), Iodo-gallic acid (**IGA**), *trans*-ferulic acid (**FA**), 2-iodo-*trans*-ferulic acid (**2IFA**) were obtained as described previously.^[20]^ Salicylic acid (**SA**; #247588), 5-iodo-salicylic acid (**5ISA**; #I10600), 3,5-diiodo-salicylic acid (**3,5diISA**; D124001), 5-iodo-2-methoxy benzoic acid (**2M,5IBA**; #754862), 4-aminosalicylic acid (**PAS**; #A79602) and acetyl-salicylic acid (**ASP**; #PHR1003) were purchased from Sigma-Aldrich (Saint-Quentin Fallavier, France) and were >95% purity. Stock solutions in which the compounds were found to be completely soluble in DMSO (DASIT group; #455103) were prepared at a 20 mg/mL final concentration (except for **ASP** which was concentrated at 10 mg/mL), prior to drug susceptibility testing.

Computed properties of all phenolic derivatives, such as molecular weight, octanol-water partition coefficient LogP, phenolic and acidic pKa, were done using the Marvin Desktop Suite, Calculator Plugins 24.1.3 developed by ChemAxon (http://www.chemaxon.com).

### Bacterial strains and culture conditions

*P. aeruginosa* PAO1 and *E. coli* ATCC 25922 were routinely grown in Mueller-Hinton II broth (MHIIB; Sigma-Aldrich; #90922) at 37 °C under agitation at 180 rpm.

*M. abscessus* CIP104536^T^ smooth (S) or rough (R) morphotype, *M. marinum* ATCC BAA-535/M, and *M. tuberculosis* ATCC 25117 H37Ra reference strains were routinely grown in Middlebrook 7H9 broth (BD Difco; #271310) supplemented with 0.2% glycerol (Euromedex; #EU3550), 0.05% Tween-80 (Sigma-Aldrich; #P1754) and 10% oleic acid, albumin, dextrose, catalase (OADC enrichment; BD Difco; #211886). *M. smegmatis* mc^2^155 reference strain was cultured in the same medium devoid of OADC supplementation. All cultures were maintained at 37 °C with shaking, except for *M. marinum* which was cultured at 32 °C.

Recombinant *M. tuberculosis* H37Ra strain expressing pH-GFP^[28]^ (pUV15-pHGFP; Addgene plasmid #70045, kindly gifted by Sabine Ehrt), was generated by electroporation according to Goude *et al*.^[37]^ and further selected onto 7H11 agar medium (BD Difco; #283810) supplemented with 10% OADC and 50 μg/mL of Hygromycin B (Toku-E; #H007). Hygromycin B was used as selection marker for the culture maintenance of the fluorescent strain at a final concentration of 50 μg/mL, but was not used when performing intrabacterial pH homeostasis disruption experiments.

### Antimicrobial susceptibility testing on Gram^(-)^ bacteria

Minimum inhibitory concentrations were determined using the rapid INT colorimetric assay.^[38]^ Briefly, fresh mid-log phase bacterial culture (OD_600_ = 0.6-0.8) was diluted to a cell density of 1 × 10^6^ CFU/mL in MHIIB. Then, 100 μL of this inoculum was added in a 96-well flat-bottom Corning® microplates with lid (Merck, #CLS3370) containing two-fold serial dilutions of each compound to a final volume of 200 μL. Growth controls containing no compound (*i.e*., bacteria only = B), inhibition controls containing standard drug (= doxycycline), and sterility controls (*i.e*., medium only) without bacteria were also included. Plates were incubated at 37 °C for 16-18 h, then 50μL of a *p*-iodonitrophenyltetrazolium violet (INT) (Sigma-Aldrich; #I8377) solution (0.2 mg/mL) was added to each well. The microplates were re-incubated in the dark at 37 °C for 30 min until the appearance of a color change in the control B-wells (bacteria alone). In the presence of active dehydrogenases, colorless INT solution is reduced to an insoluble purple formazan dye, synonymous of the presence of metabolically active bacteria. The absorbance of formazan was further measured at 470 nm with a Tecan Infinite^®^ 200 PRO multimode microplate reader (Tecan Group Ltd, France). Relative absorbance units were defined as: RAU% = (test well A_470 nm_/mean A_470 nm_ of control B wells) × 100. MIC values were determined by fitting the RAU% sigmoidal dose−response curves in Kaleidagraph 4.2 software (Synergy Software, Reading, PA). The drug concentration that caused a 90% reduction in optical density compared to the growth controls was defined as the MIC_90_. All experiments were performed independently at least three times.

### Antimycobacterial susceptibility testing using Resazurin microtiter assay (REMA)

Antimycobacterial susceptibility testing was performed using the Middlebrook 7H9 broth microdilution method. MICs were determined in 96-well flat-bottom Nunclon Delta Surface microplates with lid (Thermo-Fisher Scientific, #167008) using the resazurin microtiter assay (REMA).^[39-40]^ Briefly, log-phase bacteria were diluted to a cell density of 5 × 10^6^ CFU/mL in complete 7H9 medium. Then 100 μL of the above inoculum was added to each well containing 100 μL of complete 7H9 medium, serial two-fold dilutions of the compounds or controls, to a final volume of 200 μL (final bacterial load of 5 × 10^5^ CFU per well). Growth controls containing no inhibitor or with the DMSO vehicle (*i.e*., bacteria only), inhibition controls containing 50 μg/mL kanamycin (Euromedex; #UK0010D), and sterility controls (*i.e*., medium only) without inoculation were also included. Microplates were incubated at 37 °C (32 °C for *M. marinum*) in a humidity chamber to prevent evaporation for 3–5 days (*M. smegmatis* and *M. abscessus*) or 10-14 days (*M. marinum* and *M. tuberculosis*). Then, 20 μL of a 0.025% (*w/v*) resazurin solution in sterile water (Sigma-Aldrich; #R7017) was added to each well, and the plates were incubated at 37 °C (or 32°C) until color change from blue to pink or violet in the control well (*i.e*., bacteria alone). Fluorescence units (FU) of the metabolite resorufin (λ_ex_/λ_em_ = 530/590 nm) were quantified using a Tecan Spark 10M™ multimode microplate reader (Tecan Group Ltd, France). Relative fluorescence units (RFU) were defined as: RFU% = (test well FU/mean FU of control *B* wells) × 100. MIC values were determined by fitting the RFU% sigmoidal dose–response curves in Kaleidagraph 4.2 software. The lowest compound concentration leading to 90% inhibition of bacterial growth was defined as the MIC_90_. All experiments were performed independently at least three times.

### Cell culture and cytotoxicity assays

The cytotoxicity of compounds against eukaryotic cells was measured based on the reduction of resazurin^[22, 41]^ as a value of cellular viability by metabolic activity. Murine macrophage cell-line Raw264.7 (American Type Culture Collection TIB-71) were cultured from a cryo-preserved stock in Dulbecco’s modified Eagle medium (DMEM; Corning, #10-013-CV) supplemented with 10% heat-inactivated fetal calf serum (FBS, Sigma, #F7524) in 25 cm^2^ tissue culture flasks (Corning-Falcon; #353108). Cells were grown at 37 °C and 5% CO2 until reaching subconfluency (60-80%).

For cytotoxicity experiments, approximately 1 × 105 cells were seeded in each well of a 96-well flat-bottom Nunclon Delta Surface microplates (Thermo-Fisher Scientific; #167008) in a final volume of 200 μL, and incubated for additional 16-24 h. Then, the medium was removed by aspiration, and 200 μL of serial two-fold dilution of each compound in DMEM-FBS were added to each well. After 24 h of incubation, 20 μL of a 0.025% (*w/v*) resazurin solution was added to each well. Fluorescence was measured following incubation for ∼4 h at 37 °C and 5% CO_2_ in the dark, as described above, leading to relative metabolic activities. DMSO treated cells were used as 100% viability control conditions, and addition of 0.2% Triton X-100 solution served as positive control of total lysis (0% viability). The compound concentration leading to 50% macrophages cell death was defined as the CC_50_. All experiments were performed in technical triplicates on two independent assays.

### pH-dependent growth inhibition assay and MIC determination

Susceptibility testing was performed as described above with slight modifications. All experiments at pH 6.8 were performed in standard Middlebrook 7H9 containing 0.2% glycerol and 10% OADC enrichment. To avoid Tween-80-mediated toxicity, 0.025% Tyloxapol (Sigma Aldrich; #T8761) was used. For pH-dependent experiments the same media containing additional 50 mM of MES and adjusted at pH 5.5^[27, 42]^ was used. The drug of interest was two-fold serial diluted in the appropriate media to a final volume of 100μL per well in technical triplicate per microplate. Then, each well was inoculated with 100μL (5 x 10^6^ CFU/mL) of a *M. tuberculosis* culture in pH 6.8 or pH 5.5-ajusted Middlebrook 7H9. Positive growth controls (*i.e*., inoculum without antibiotics or with DMSO), positive growth inhibition controls (*i.e*., 50 μg/mL kanamycin), and sterility controls (*i.e*., medium only) were included. The 96-well flat-bottom Nunclon Delta Surface microplates were incubated at 37 °C during 14-21 days for both pH 6.8 and pH 5.5 experiments. MIC of each antibiotic at each pH was determined by visual inspection according to EUCAST recommendations^[43]^, and by absorbance reading at 600 nm to confirm visual recording. The first antibiotic concentration that visually inhibited bacterial growth was defined as the MIC_90_.

### Intrabacterial pH homeostasis disruption assays

Recombinant *M. tuberculosis* harboring pUV15-pHGFP and producing the ratiometric pH-sensitive pH-GFP sensor was used for the determination of intrabacterial pH homeostasis disruption.^[28, 32-33]^ Briefly, exponentially growing *M. tuberculosis* cultures were centrifuged at 3,500 rpm during 5 min, and resuspended in 10 mL of Middlebrook 7H9 adjusted at pH 5.5 or pH 6.8 in order to have a normalized OD_600nm_ of 0.8. Then, 100μL of this bacterial inoculum was used to inoculated wells containing 100 μL of serial-diluted compounds of interest, giving a final OD_600nm_ of 0.4 in each well. Prior performing the IBpH perturbation experiments, we have checked that all the compounds, at their highest concentration tested, did not impact the pH of the medium by more than 0.1 pH unit. The pH-GFP fluorescence was collected at λ_em_ 535 nm after excitation at λ_ex_ 405 nm and λ_ex_ 488 nm, respectively, using a TECAN Spark 10M™ multimode microplate reader (Tecan Group Ltd, France). For all our experiments at pH 5.5, the well-established ionophore Monensin (Sigma; #M5273) was used at 20 μM as internal positive control of intrabacterial pH homeostasis disruption. For all our experiments at pH 6.8, the well-established protonophore CCCP (Merck; #215911) was used at 100 μM as internal positive control of intrabacterial pH homeostasis disruption. The ratios in fluorescence intensity λ_ex_ 405 nm / λ_ex_ 488 nm from each condition were calculated. All the results were exported as CSV files, imported in the R Studio software

(The R Project for Statistical Computing, version 1.3.1073), and graphs were plotted with the ggplot2 package (version 3.3.2). Determination of half-maximal effective concentration (EC_50_) was performed in the R Studio software using a four-parameter logistic nonlinear regression model.^[33]^ Finally, ΔRatio_pH6.8_ or ΔRatio_pH5.5_ were determined by subtracting the minimal fluorescence ratio values from their respective maximal fluorescence ratio values obtained in the fitted model.

### Spontaneous resistant mutant isolation

Approximately, 10^6^ CFU of exponentially growing *M. tuberculosis* H37Ra were plated on a single Petri dish of Middlebrook 7H11 agar medium supplemented with 10% OADC and increasing concentrations of **SA, 5ISA, 3,5diISA, ASP** or **PAS** respectively (corresponding to 1×, 2× or 5× MIC_90_ at pH 6.8). Additional plates without compounds were included in the experiments as growth control. Plates were incubated at 37 °C and a thorough visual inspection of resistant clones’ appearance was done after 4 and 6 weeks. Experiments were performed at least on two independent biological replicates.

### Quantification and statistical analysis

All results displayed in this study were obtained from *n* = 2 or *n* = 3 biologically independent experiments performed at least each time in three technical replicates (unless otherwise stated). For statistical analysis in the intrabacterial pH homeostasis disruption assays, the means between the conditions of interest were tested for significant differences using one-way ANOVA followed by Tukey’s post-test with the ‘aov()’ and ‘TukeyHSD()’ functions in R – R Studio using the ggpubr R package. All the *p*-values contained in the text or the figures are relative to the control condition (unless otherwise stated). All *p*-values were considered significant when *p-value* < 0.05. Statistical analysis is displayed in the figures as: n.s, not significant; *p*-value < 0.05; *p*-value < 0.01; or *p*-value < 0.001. In each figure or table legend, the statistical tests used, the number of biologically independent replicates and the number of technical replicates is indicated.

## Abbreviations

2IFA: 2-iodo-ferulic acid
2M,5IBA: 5-iodo-2-methoxy benzoic acid
3,5diISA: 3,5-diiodo-salicylic acid
5ISA: 5-iodo-salicylic acid
ASP: aspirin
BA: benzoic acid
CC_50_: Cytotoxic concentrations leading to 50% macrophages cell death
GA: Gallic acid
IBpH: intrabacterial pH
IGA: 2-iodo-gallic acid
MIC_90_: Minimum inhibitory concentration leading to 90% bacterial growth inhibition
PA: Phenolic acids
PAS: 4-amino-salicylic acid / *para*-aminosalicylic acid
SA: Salicylic acid
TB: Tuberculosis
*t*FA: *trans*-ferulic acid

## Acknowledgements

We would like to acknowledge all members of the Lipolysis and Bacterial Pathogenicity group and the LISM unit for continuous support and insightful discussions. This work was supported by the Centre National de la Recherche Scientifique (CNRS) and Aix-Marseille Université (AMU). PS received financial support from the CNRS Biologie, the Agence Nationale de Recherches sur le Sida et les Hépatites virales (ANRS) (project n°ANRS0358) and the French government under the France 2030 investment plan, as part of the Initiative d’Excellence d’Aix-Marseille Université - A*MIDEX and is part of the Institute of Microbiology, Bioenergies and Biotechnology - IM2B (AMX-19-IET-006). PS has also received a FEBS Excellence Award and would like to deeply thank the FEBS Fellowships Office for its continuous support. J.L., T.F. and E.F. PhD fellowships were supported by the Ministère de l’Enseignement Supérieur et de la Recherche Française.

The funders did not play a role in the study design, data collection and analysis, decision to publish, or preparation of the manuscript.

## Data availability statement

Any additional data that support the findings of this study are available upon reasonable request from the corresponding authors at.

## Table of Contents graphic

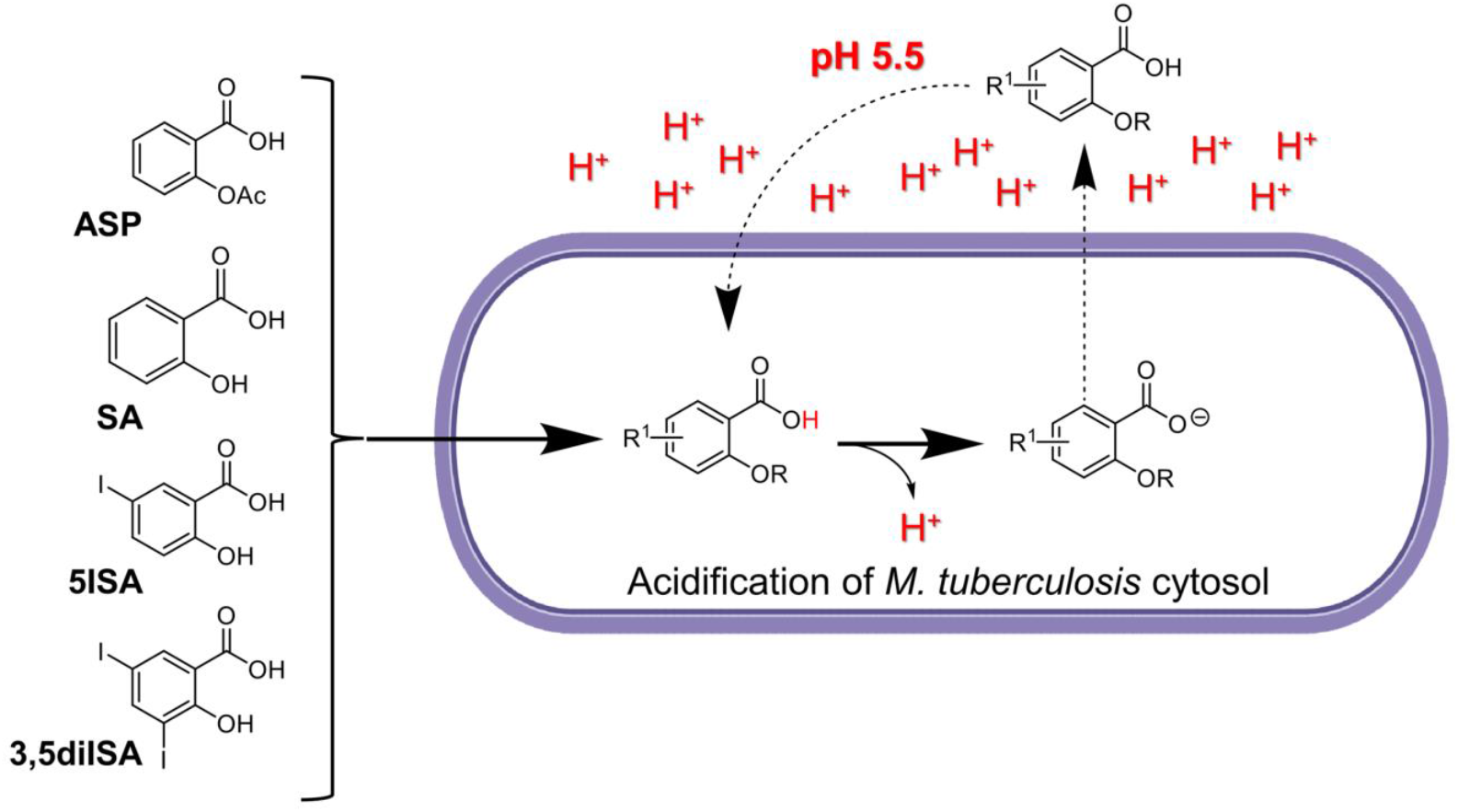

Salicylic acid (**SA**), its two iodinated analogues **5ISA** and **3,5diISA**, as well as the anti-inflammatory drug aspirin (**ASP**), are anti-TB drugs potentialized at acidic pH which act by disrupting the intrabacterial pH of *M. tuberculosis* in a dose-dependent manner, under *in vitro* conditions mimicking the endolysosomal pH of macrophages.

